# *Bloom* unifies brain cell representations across modalities and contexts

**DOI:** 10.64898/2026.11.11.765881

**Authors:** Rohan Gala, Ian Convy, Yeganeh Marghi, Ruiyang Liu, Jeremy Roghair, Renhao Xue, Nathan Gouwens, Sven Otto, Samantha Caldwell, Krutika Khinvasara, Jon Martin, Sudipta Sengupta, Zizhen Yao, Michael Hawrylycz, Shoaib Mufti, Hongkui Zeng, Uygar Sümbül

**Affiliations:** Allen Institute, Brain Science, Seattle WA, USA; Amazon Web Services, Seattle WA, USA

## Abstract

Multimodal genomic datasets now profile mammalian brain cells across anatomical regions, developmental stages, disease states, and species. Learning shared representations that capture cellular diversity across these heterogeneous observations remains a central challenge. Here, we introduce Bloom, a transformer-based model that represents cells using gene expression, chromatin accessibility, or both. Bloom is trained on 300,000 adult mouse brain cells with paired measurements, randomly sampled with broad-subclass balancing—approximately 100-fold fewer cells than used by comparable pre-trained models. By combining genomic sequence embeddings with randomly sampled multimodal cell views, Bloom outperforms unimodal cell models in resolving brain-wide cell types and in annotation, regression, and label-transfer tasks. Its attention patterns reveal cell-type-dependent associations among genomic loci, including pairs separated by millions of base pairs. These results establish Bloom as a data-efficient framework for integrating multimodal brain cell measurements and generating hypotheses about cell-type-specific genomic relationships.

## Introduction

Cellular phenotypes emerge from complex relationships among molecular and functional features. Gene expression and chromatin accessibility, for example, are intricately linked,^1^ yet technologies that measure these features introduce distinct biases. Multimodal characterization is therefore essential for resolving these relationships and motivates a shared representation of cells that integrates observations across modalities.^2, 3^ This need is particularly acute in the brain, where immense cellular diversity^4^ remains characterized through largely fragmented measurements.^5^

The flexibility of attention mechanisms^6^ allows cells to be encoded from different feature sets, including non-identical gene sets. Yet even high-dimensional embeddings learned from large datasets can substantially distort the underlying cellular geometry, despite both large embedding dimensionality and training corpus.^7, 8, 9, 10^ These distortions may partly reflect reliance on RNA sequencing ^11, 12, 13, 14, 15, 16, 17, 18^ and selected gene sets, such as highly variable genes, which introduce technology- and feature-selection biases. Furthermore, pre-trained models often separate cells in latent space according to the profiling technology (e.g. single-cell RNA sequencing (scRNA-seq), single-nucleus RNA sequencing (snRNA-seq), and spatial transcriptomics) rather than aligning biologically corresponding populations. This separation limits the benefits of training on millions of cells from diverse sources and undermines a central goal: a shared representation that supports cell identification and comparison across experimental platforms.

Here, we introduce Bloom, a multimodal^19^ pre-trained cell model focused on brain cells and genomic measurements. During training, Bloom receives randomly sampled views of each cell comprising gene expression, chromatin accessibility, or both, drawn from a feature library of over 32,000 genes and 1.7 million accessibility peaks. Although we focus on genomic modalities because of data availability, Bloom’s architecture also accommodates non-genomic features, including spatial location, dendritic and axonal morphology, connectivity, and physiology. Previous models have represented genes using arbitrary identifiers ^11, 12, 13^ or embeddings from protein language models,^20, 14, 15^ which do not directly represent non-coding DNA. In contrast, Bloom combines embeddings from genomic language models ^21, 22^ with relative chromosomal position to tokenize genomic sites independently of the observation modality. This approach provides a shared framework for representing coding and non-coding regions across modalities and supports training and inference across species.

The scaling gains that have driven advances in natural language models have remained largely elusive for tabular single-cell data.^23^ One possible explanation is that, when inputs are restricted to selected genes, additional cells of the same type and similar biological background provide limited new information. Bloom achieves state-of-the-art performance in classification, clustering, label transfer, and alignment across mouse brain gene expression and chromatin accessibility datasets, despite training on approximately 100-fold fewer cells than existing pre-trained models. The training dataset requires minimal curation, with balancing limited to capping the number of cells per broad subclass. This efficiency may partly reflect Bloom’s random sampling of cell views from approximately 1.8 million multimodal features, which provides a form of data augmentation. Bloom’s representations are correspondingly robust to variation in both the number and identity of input features.

Finally, we examine Bloom’s first attention layer to investigate gene–peak associations. The model learns statistically significant, cell-type-dependent associations whose strengths show little dependence on either genomic distance or the fraction of cells in which the gene is expressed and the peak is accessible. Together, these advances distinguish Bloom through a combination of modality-independent genomic tokenization and learning from randomly sampled multimodal cell views. Bloom expands the diversity of observations through which each cell is represented. Its architecture also provides a foundation for incorporating additional cellular modalities and extending shared representations across species.

## Results

### Bloom architecture and multimodal tokenization

To enable shared representations of cells across modalities, we developed Bloom, a single-cell multimodal transformer-based model. As a multimodal model, Bloom accesses a large and diverse set of cell features, including coding and non-coding sequences, while maintaining parameter and data efficiency (**Figure 1**). Recent models represent genes using embeddings of their encoded amino acid sequences,^20, 14, 15, 16^ an approach that does not extend directly to non-coding RNAs or chromatin accessibility peaks (**Figure 1**). Bloom instead represents genomic features using embeddings of their underlying DNA sequences, which can capture sequence patterns relevant to gene regulation and gene–peak relationships. This approach accommodates both coding and non-coding regions, substantially expanding the feature library within a shared representation framework (**Figure 1**; **Methods**).

**Figure 1.**
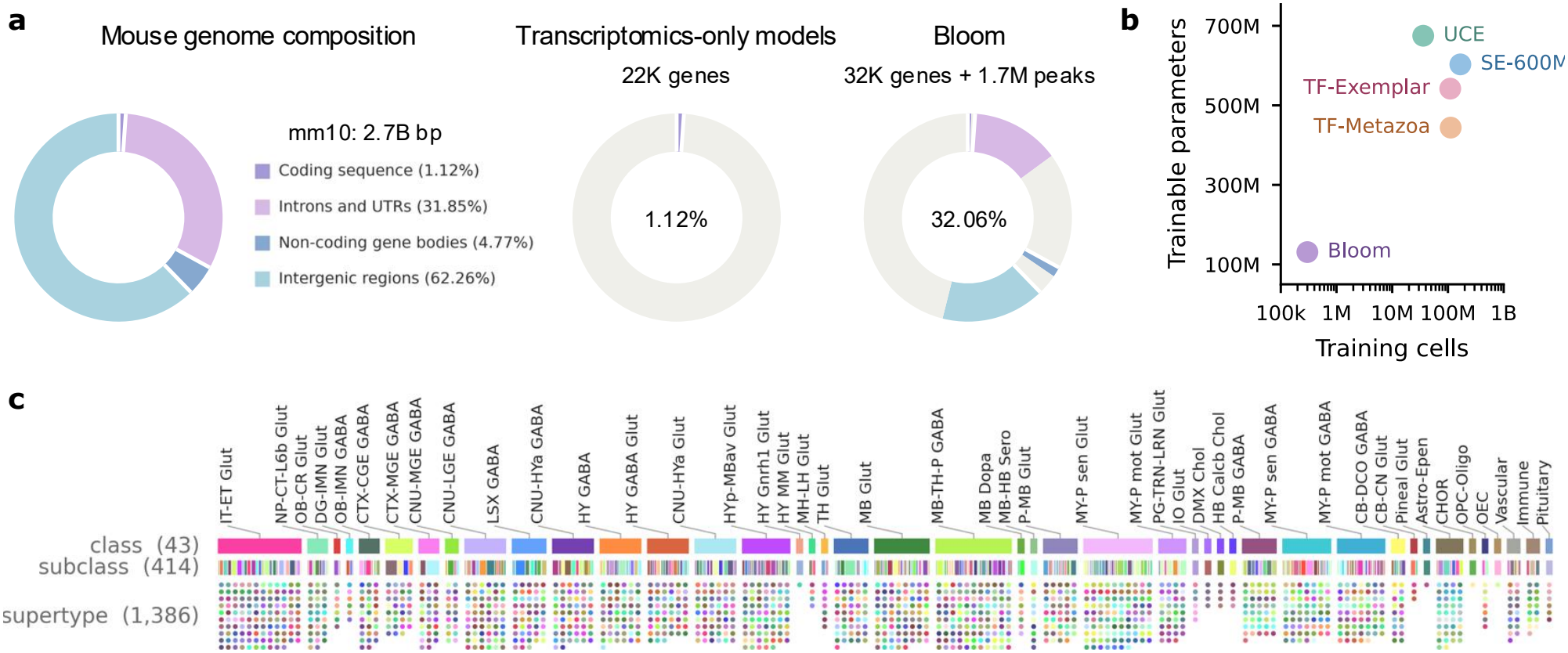
Bloom explores a diverse fraction of the mouse genomic landscape while maintaining high parameter and data efficiency. **a)** Mouse genome composition and the fraction of genomic base pairs covered by the input features of different modeling approaches (Methods). Unlike transcriptomic models, which restrict their input feature space to only protein-coding sequences, Bloom incorporates a significantly broader genomic landscape. Its input feature space spans coding and non-coding sequences, as well as introns, untranslated regions (UTRs), and intergenic regions, capturing a more holistic view of the genome. **b)** Comparison of model scale and dataset size in terms of parameter count and number of cells used to train prominent single-cell foundation models (Table 3). Bloom consists of about 130M trainable parameters while being trained on a multimodal dataset of 300K cells. In contrast, existing models such as UCE, State (SE-600M), and Transcriptformer are substantially larger architectures (400M–700M parameters) that are trained on datasets that are orders of magnitude larger in terms of number of cells. **c)** The Allen Institute mouse whole-brain reference taxonomy^4^ consists of hierarchically organized labels, including 43 classes, 414 subclasses, and 1,386 supertypes in the current version (AIT33).

For the whole mouse brain single-cell RNA sequencing and chromatin accessibility (multiome) datasets analyzed here, this corresponds to a feature set of 32,129 genes and 1,747,701 peaks. While genes are well-defined in reference genomes, peaks are defined as regions of 500 base pairs (bp) along the genome based on pooled ATAC-seq data (**Methods**). Thus, Bloom randomly samples cell views from a set that is ∼2 orders of magnitude larger than models considering only the transcriptome (**Figure 1**).

Each genomic feature is encoded as a combination of its underlying DNA sequence, its genomic location, and its measured value in a given cell (**Figure 2a**), allowing Bloom to tokenize genomic sites independently of the observation modality. Rather than collapsing these components into a single embedding, Bloom’s encoder uses custom attention heads that attend over them separately; sequence heads compute queries and keys from the DNA-sequence embedding, position heads from the genomic-location embedding, and standard heads from the full token (**Figure 2b**). To efficiently process these long sequences, the resulting tokens are passed to a *Perceiver*^24^ block.

**Figure 2.**
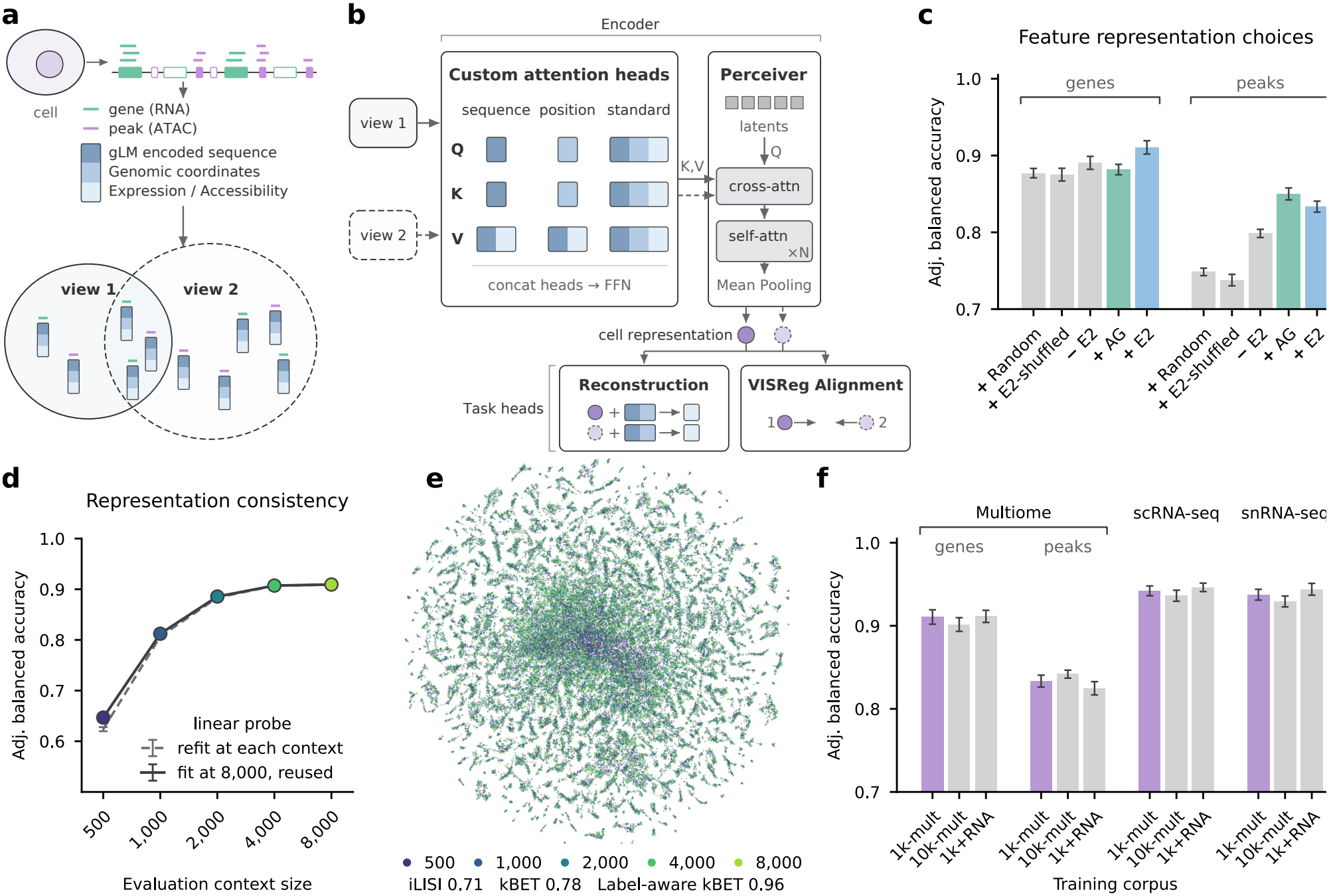
Bloom architecture and scaling. **a)** Feature encodings for multimodal data. Each gene or peak feature is represented by its gLM-encoded DNA subsequence, genomic location, and measurement in a given cell. Views of a cell are constructed by sampling features from among expressed genes or accessible peaks in that cell. **b)** Tokenization to cell embedding, with reconstruction and alignment objectives. Sequence and position attention heads compute key, query, and value vectors from a subset of token components. Outputs from the different heads are concatenated and pass through a feed-forward layer (FFN). The resulting tokens are compressed by Perceiver cross-attention into a fixed set of learned latents, refined by *N* self-attention layers and mean-pooled into one cell embedding per view (solid, view 1; dashed, view 2), which feeds two task heads: reconstruction of query features and alignment between views. **c)** Effect of sequence and position encodings on downstream classification performance. Cell type classification with Multiome dataset cell representations obtained using gene-only, or peak-only inputs are considered as separate evaluation tasks. While performance improves modestly with gLM-derived sequence encodings over randomly initialized encodings on the gene-only task, the advantage is substantial on the peak-only task. E2: Evo 2; AG: AlphaGenome; E2-shuffled: Evo 2 sequence embeddings shuffled across features. +/*−* prefix: with/without position encodings. **d)** Bloom representation robustness to variations in input context length. The solid line shows accuracy of a linear classifier trained on representations obtained with a context size of 8,000 genes and reused at the smaller context sizes. The dashed line shows performance of classifiers that were trained for each context size independently (Methods). The minimal gap between the two curves indicates that a single classifier can be robustly reused across different input sizes and suggests that representations degrade gracefully with reduced context lengths. **e)** Joint UMAP projection of representations for the same cells obtained with different input context sizes (colors). Overlap of colors suggests cell representations are well-integrated across input context sizes. High iLISI, kBET, and label-aware kBET (subclass labels) metrics quantify this aspect. **f)** Impact of training dataset scale and composition on downstream classification accuracy across Multiome and RNA-only datasets. Datasets were balanced to a maximum of 1K cells per subclass (`1k-mult`), 10K cells per subclass (`10k-mult`), or supplemented with additional unimodal RNA data (`1k+RNA`). Expanding the training corpora, either by increasing the subclass sampling limit (`10k-mult`) or by incorporating unpaired unimodal data (`1k+RNA`), does not yield substantial performance improvements over the baseline `1k-mult` configuration. Error bars in **c, d**, and **f** show *±*1 SEM over 10 cross-validation folds.

Bloom is trained with two complementary objectives on top of the cell embedding (**Figure 2b**). For the reconstruction objective, we use a fixed set of ∼8,000 differentially-expressed genes as targets for all cells. The decoder head conditions on the cell embedding together with a gene’s sequence to predict whether the gene is present or absent in the cell (**Methods**). For the alignment objective, a separate head applies VISReg^25^ regularization to the embeddings of two views of the same cell, encouraging the latent space to capture cell state while remaining invariant to view-specific variation, including the modality used to profile the cell (**Figure 2b**; **Methods**).

### Unbiased cell views sampled from multimodal features augment data

Considering that existing sequencing methods are significantly biased towards non-expression (i.e. gene/peak dropout),^26^ Bloom samples only the non-zero genes and peaks of each cell (**Methods**). In the whole mouse brain multiome corpus we used for training, each cell expresses ∼8K genes and has ∼40K accessible peaks on average. Moreover, considering that different sc/snRNA-seq or multiome platforms display significant domain shifts in both the dropout noise and the expression values (e.g., sequencing from soma vs nucleus, differences in capture rates and amplification, errors in aligning measurements to the reference genome), Bloom binarizes each genomic measurement to minimize related batch effects (**Methods**).

At each iteration, Bloom samples a batch of cells and forms two independently sampled views of each cell. Bloom performs gradient descent to jointly minimize the discrepancy between the two representations corresponding to the same cell^25^ and the error in predicting binarized expression of the query genes (**Figure 2a–b**; **Methods**).

### Expanded genome featurization by DNA sequence- and location-based tokenization

We evaluated how genomic sequence representations contribute to Bloom’s performance by comparing embeddings from two recent genomic language models, Evo 2 ^21^ and AlphaGenome,^22^ with randomly assigned feature embeddings. Performance was measured by the accuracy of a linear classifier trained on Bloom’s cell embeddings to predict subclass identity. Evo 2 and AlphaGenome yielded comparable overall accuracy, with Evo 2 performing better for gene-based inputs and AlphaGenome for peak-based inputs (**Figure 2c**). Both outperformed random feature embeddings, demonstrating that genomic sequence information improves cell representations beyond what Bloom can learn from arbitrary feature identifiers. This advantage was greater for accessibility peaks, whose feature library is substantially larger than that of genes. Shuffling Evo 2 embeddings across features reduced accuracy to a level comparable to random embeddings (**Figure 2c**), indicating that the improvement depends on matching sequence embeddings to the correct genomic features, rather than on their marginal statistical properties. These results support the value of sequence-derived information for learning cellular representations, although they do not establish which sequence properties, such as transcription factor binding motifs, drive the improvement.

Genomic position provides additional context for relationships among genes and accessibility peaks. Bloom therefore incorporates chromosomal position during tokenization, using relative rather than absolute coordinates to avoid dependence on a particular reference genome’s coordinate system and support application across species (**Methods**). Although previous work found no benefit from absolute positional information in in-distribution evaluations,^14^ relative positional encoding improved Bloom’s performance compared with omitting position information (**Figure 2c**; **Methods**). Thus, sequence and relative genomic position provide complementary information for representing cells.

Bloom also uses a context length that is four times that of SE-600M and Transcriptformer, and eight times that of UCE (**Table 3**), allowing more features to be considered jointly when constructing a cell representation. Increasing the context length improved subclass classification accuracy (**Figure 2d**). During training, Bloom randomly samples both the identity and number of input features, up to the maximum context length, exposing the model to views ranging from sparse feature panels to broader genomic profiles. This strategy addresses the variable feature coverage of experimental platforms: Bloom’s representations maintain robust classification performance across a wide range of input feature counts (**Figure 2d**).

### A multimodal encoder-only Transformer model

Different technologies and experiments characterize cells based on different sets of features. Therefore, inferring consistent representations based on unequal numbers of (and perhaps even non-overlapping) observations is a key methodological need. The degree of such consistency across observation modalities is itself an open problem for brain cells. Moreover, this methodological flexibility could enable *in silico* experiments (e.g., what happens if a peak becomes accessible), thereby hastening new biological insights. Bloom attempts to learn such representations with an encoder-only Transformer model.^27^ Transformers can handle unequal input lengths gracefully via their attention mechanism, and can be designed to yield permutation invariance with respect to their inputs.^28, 29^ Finally, useful relationships of tabular features can be captured or provided via feature-wise embeddings,^30^ as described above. Previous approaches to model single-cell transcriptomic data^14, 15^ have used protein foundation models^31^ to *tokenize* genes. Bloom expands this approach to multiple modalities via its universal tokenization, unbiased cell view sampling, and a self-supervised loss to align different cell views.

Bloom also expands to include non-coding features while supporting multiple species via its attention to relative distances between subsequences on the chromosome and its use of pre-trained DNA sequence embeddings.^32, 33, 21, 22^ For features that are on different chromosomes, Bloom’s location-based attention is near zero by construction, but it can still learn feature associations based on the relationship between the underlying DNA sequences (**Figure 2b**; **Methods**). We validate that DNA sequence embeddings enable non-trivial performance on tasks defined over gene and peak subsequences relevant for our data (e.g., predicting overlap of a given sequence with known promoter regions, UTR sequences, satellite regions). Noting that the performance on these tasks doesn’t drop substantially if the embeddings undergo linear dimensionality reduction with PCA, we use reduced dimensionality embeddings to reference gene and peak features in Bloom.

### Bloom achieves state-of-the-art performance in transcriptomic classification of mouse brain cells

Accuracy in assigning correct cell type identities to cells based on existing taxonomies is used as a standard metric in judging representation quality, which is performed with linear classifiers to avoid the warping introduced by more powerful classifiers.^40^ At the subclass level across the whole adult mouse brain (**Table 2**), Bloom achieves 91% (multiome, 315 subclasses), 94% (scRNA-seq,^4^ 310 subclasses), and 94% (snRNA-seq,^34^ 297 subclasses) cross-validated adjusted balanced accuracy in linearly classifying brain cells based on gene expression although it was trained only with multiome data and improves upon the baseline methods considered here (**Figure 3a,c,d**). Bloom achieves these accuracies while *jointly* learning to represent cells based on chromatin accessibility information. The other models consider only gene expression. Hence, only Bloom appears twice in **Figure 3a**. Interestingly, Bloom’s subclass accuracy based on chromatin accessibility is better than that for the Transcriptformer and UCE models based on gene expression, even though single-cell accessibility measurements are considered more heterogeneous within cell types.^41^

**Figure 3.**
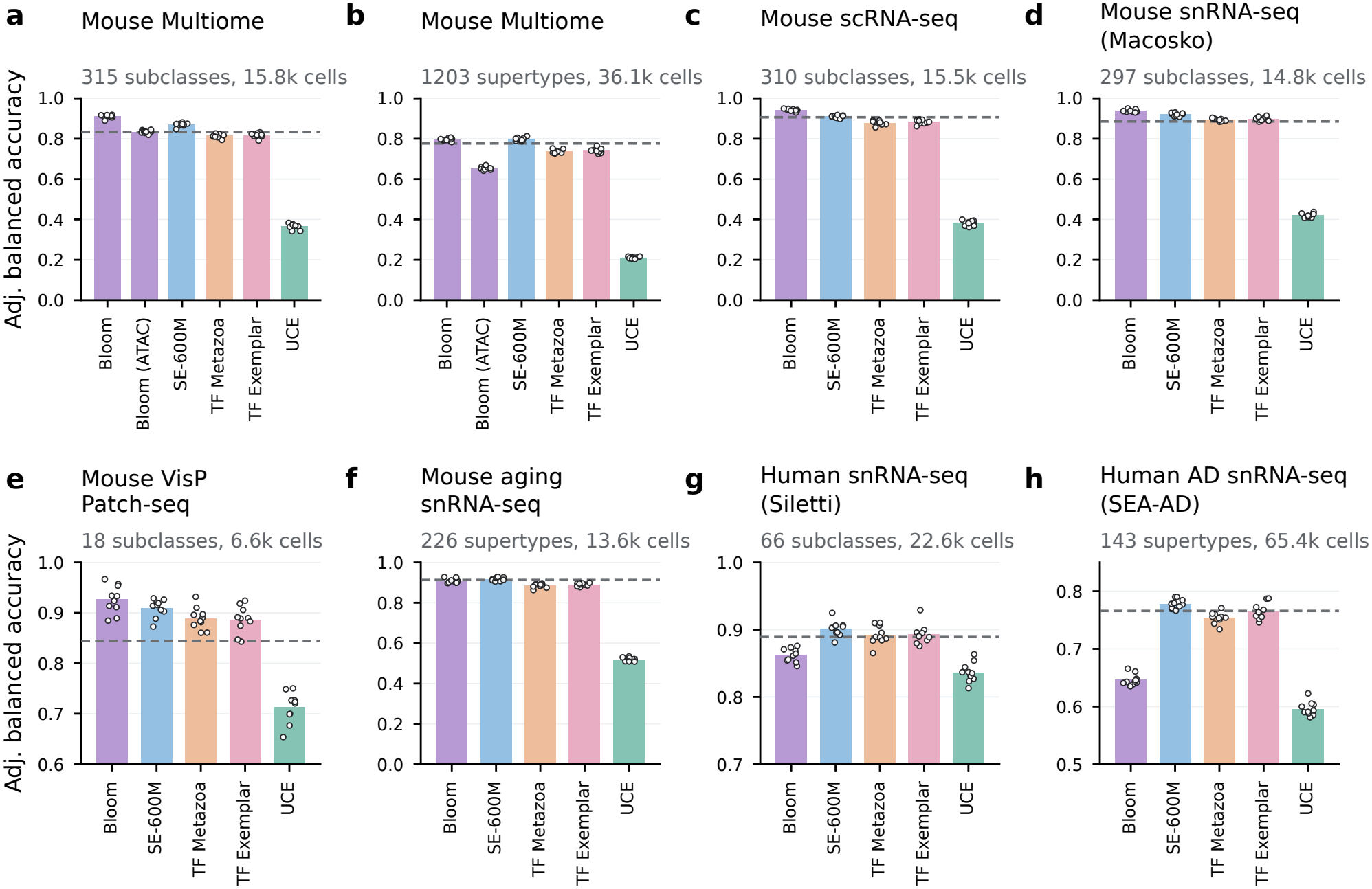
Bloom achieves state-of-the-art results in the classification of mouse brain cells. Performance of linear classifiers on held-out evaluation datasets using representations obtained from different models (Methods). Bars show mean adjusted balanced accuracy across folds; markers show individual folds. Dashed lines provide a baseline calculated with 1,024-component gene-only PCA representations. All results are based on gene-only inputs, except for *Bloom (ATAC)*, where the cell representations are obtained with peak-only inputs. Bloom is trained on only adult mouse brain Multiome data (`1k-mult`). Datasets in **(a–h)** are increasingly out-of-distribution in relation to the Bloom training corpus. **(a)** Mouse brain Multiome dataset, subclass labels. **(b)** Mouse brain Multiome dataset, supertype labels are finer resolution than subclass labels. **(c, d)** Mouse brain scRNA-seq^4^ and snRNA-seq^34^ datasets. **(e)** Mouse VisP Patch-seq.^35, 36^ Patch-seq uses a version of SMART-Seq for transcriptomic profiling. **(f)** Mouse brain aging snRNA-seq.^37^ Datasets in **(a–e)** are from the adult mouse brain. **(g, h)** Human snRNA-seq^38^ and Alzheimer’s disease snRNA-seq (SEA-AD),^39^ to assess cross-species transfer. Note that vertical axes in **e, g**, and **h** do not start at 0.

Bloom’s performance on the scRNA-seq and Macosko snRNA-seq datasets (**Figure 3c,d; Table 2**), which were not used during training, prompts us to test its utility for datasets that can be considered progressively more different (“out of distribution”). Bloom is the top performer in the mouse Patch-seq dataset^35, 36^ and is tied for the top with SE-600M in the aging mouse dataset^37^ (**Figure 3e,f**). Despite not seeing any human data, Bloom’s performance in the human Siletti^38^ and Alzheimer’s disease (SEA-AD^39^) datasets (**Table 2; Methods**) is better than UCE’s and trails SE-600M, TF-Metazoa, and TF-Exemplar by less than 5% on the populations in healthy brains (**Figure 3g,h**). All four of these recent pre-trained baselines include millions of human cells in their training corpus. The lower performance of Bloom on the SEA-AD dataset may be due to the highly similar and continuous nature of supertypes in the human cortical areas.

Bloom is ∼5 times smaller in terms of parameter count and trained with 2–3 orders of magnitude fewer cells compared to the baselines considered here (**Figure 1b**). We find that naively increasing the number of samples in the training corpus by an order of magnitude by adding new multiome cells of the same kind does not further improve accuracy (**Figure 2f**), supporting previous observations that simply adding more cells may not lead to favorable scaling.^23^ We also find that adding unimodal snRNA-seq cells does not improve performance either (**Figure 2f**). Details of the training dataset are provided in **Table 1**.

**Table 1.** Training corpora used to train Bloom models. ^†^Multiome data from the Allen Institute (unpublished). The subclass count for 1k+RNA is the union of subclass labels across its three components.

| Corpus | Composition | Species | *n* (cells) | *n* (types) | Notes |
|---|---|---|---|---|---|
| 1k-mult | Multiome<sup>†</sup> | Mouse | 292,221 | 307 | Paired snRNA-seq and snATAC-seq; ≤ 1,000 cells per subclass (min. 500). |
| 10k-mult | Multiome<sup>†</sup> | Mouse | 1,249,786 | 307 | Paired snRNA-seq and snATAC-seq; ≤ 10,000 cells per subclass (min. 500). |
| 1k+RNA | Multiome<sup>†</sup> + scRNA-seq<sup>4</sup> + snRNA-seq (Macosko)<sup>34</sup> | Mouse | 861,429 | 351 | 1k-mult (292,221) + scRNA-seq, 10x v2 and v3 (295,149) + snRNA-seq (Macosko) (274,059); ≤ 1,000 cells per subclass per component (min. 500). |

### Bloom representations capture the underlying landscape

We next evaluate the unsupervised grouping ability of different models. We quantify the Adjusted Rand Index (ARI), Adjusted Mutual Information (AMI), and Normalized Mutual Information (NMI) values for subclass labels on three datasets. Across the three metrics and the three datasets, we find that Bloom offers substantial improvements in the clustering accuracy of cells (**Figure 4a,b,c**). Interestingly, the clustering scores of the SE-600M model, which produced competitive results in the classification study (**Figure 3**), are similar to those of a simple, dimensionality-matched linear baseline whose representations are the loadings of top principal components for each cell. The scores of the TF-Metazoa, TF-Exemplar, and UCE models are also comparable to those of the linear baseline.

**Figure 4.**
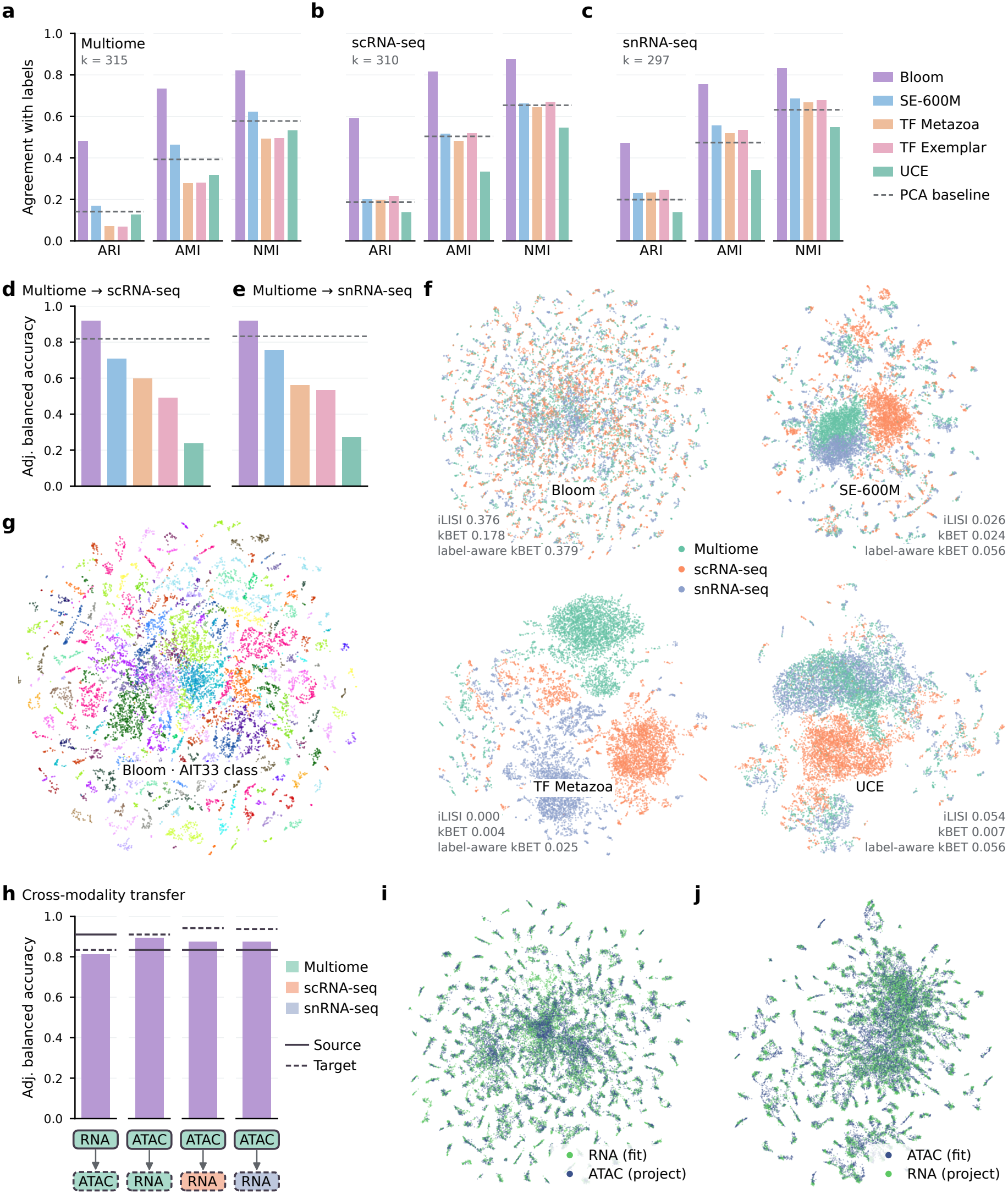
Bloom representations preserve the landscape of brain cells better, enable label transfer across measurement technologies, and are consistent across modalities. **(a–c)** K-means clustering of cell representations from three transcriptomic datasets, with *k* set to the number of reference subclasses. Agreement of clusters with subclass labels is quantified with adjusted Rand index (ARI), adjusted mutual information (AMI), and normalized mutual information (NMI). **(d, e)** Within-modality label transfer. A classifier trained on Multiome dataset cell representations is used to predict labels for scRNA-seq (**d**) and snRNA-seq (Macosko) (**e**) datasets. Dashed lines in (a–e) indicate baseline performance with 1,024-component PCA representations. **(f)** A joint UMAP for three distinct datasets using representations obtained with different models. Higher iLISI, kBET, and subclass label-aware kBET values indicate better cross-dataset integration. **(g)** Same as **(f)** for Bloom representations, but colored by class labels. Colors for distinct classes appear consistently across datasets. **(h)** Cross-modality label transfer. Classifiers trained on a source modality/dataset are used to predict labels for cells in a target modality/dataset. Solid and dashed lines on the plot indicate within-modality reference classifier performance (**Figure 3a, c, d**). **(i, j)** UMAP fit on representations of RNA-only (**i**) or ATAC-only (**j**) inputs, applied to representations of the other modality.

### Bloom representations form a common framework across profiling technologies

A main utility of large, pre-trained models is their consistent representation across technologies,^42^ enabling transfer of insights and cross-identification of cells in different studies. However, due to their unsupervised objectives (e.g., masked regression), batch effect removal cannot emerge in the absence of a relevant inductive bias. That is, if the model learns to predict some of the features given others, it will also learn to encode the batch information to improve reconstruction accuracy. As described above, Bloom binarizes the genomic features, considering that the count values depend on the underlying technology. While the level of expression of a gene carries information beyond it being expressed, we reason that Bloom’s large context size (thousands of features per cell view) can make up for the decreased information per feature. Indeed, Bloom produces high-quality cell representations (**Figure 3, 4a,b,c**). Hence, we test whether binarization has the desired effect of harmonizing over different technologies. We find that Bloom offers substantially higher accuracies in transferring labels to scRNA-seq and snRNA-seq datasets compared to baseline methods (**Figure 4d,e**; **Methods**). Unlike the baseline methods, Bloom’s training corpus does not include any cells profiled by these technologies. **Figure 4f** provides a visualization of this observation where, unlike baseline methods, Bloom embeddings mix strongly across technologies.

We also consider transferring labels from representations obtained based on gene expression alone to those obtained with peak accessibility alone. Bloom is uniquely able to perform this task among the benchmark models and is trained explicitly to accomplish this via its self-supervised training objective that aligns different views of the same cell. Indeed, we find that the accuracy of this cross-modal transfer task approaches that of within-modality predictions (**Figure 4h**) and representations based on the two modalities strongly overlap in UMAP plots (**Figure 4i,j**).

### Bloom as a hypothesis generation tool

The vastness of the search space in discovering interactions on the chromosome is a key challenge in genome biology. Bloom learns associations across millions of genetic loci, including loci that are millions of base pairs apart. These associations can potentially be queried in different ways to address a large set of questions. Here, we provide case studies to mine these learned associations.

We first characterize the distribution of gene–peak attention weights in the training dataset (i.e., `1k-mult` in **Table 1**). We choose random views for each cell in the dataset and calculate the average weight of each gene–peak pair over the sequence attention heads for that cell view (**Methods**). This produces 962 billion weight values. We consider these as the “background” and display their histogram in **Figure 5a**. The median of the attention values calculated for gene–gene pairs is slightly higher, corresponding to the ∼56^th^ percentile of the background, consistent with scRNA-seq being less noisy than scATAC-seq. Gene–peak pairs on the same chromosome and less than 1 Mbp away from each other (proximal cis pairs) receive slightly more attention, median at the ∼61^st^ percentile of the background, *n* = 23,715,315. We next study the enhancer pairs identified in Ref.^43^ for the cortex. When calculated over the enhancers’ target subclasses, these enhancers receive a median attention at the ∼81^st^ percentile of the background, *n* = 127.

**Figure 5.**
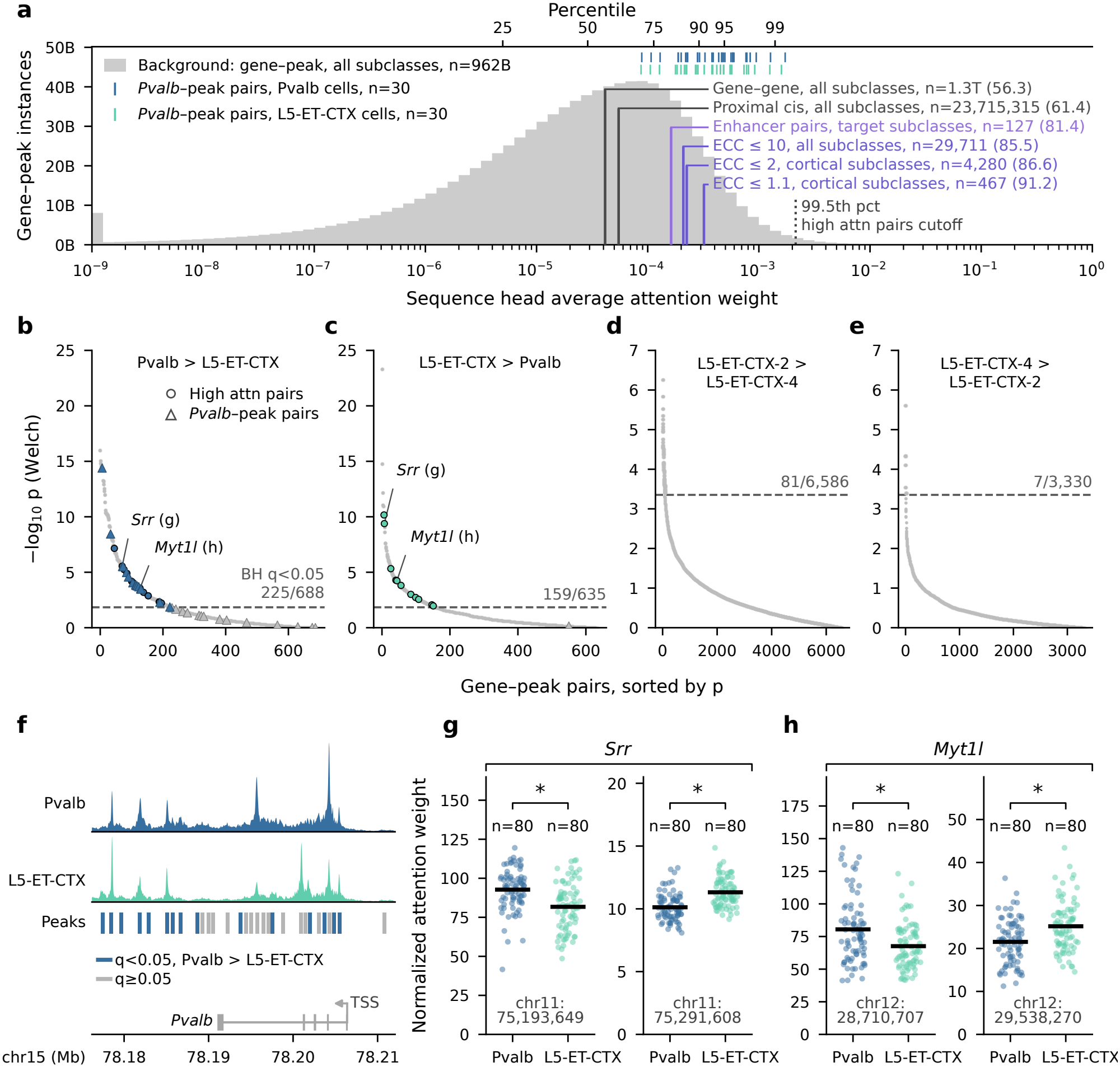
Bloom generates hypotheses for context-relevant gene–peak interactions. **a)** Attention weights for different biologically meaningful feature pairs are higher than those for background pairs (Methods). The percentiles of the median values for these feature pair sets relative to the background distribution are shown in parentheses. Attention weights are systematically larger for gene–peak pairs that are co-present in fewer subclasses (smaller ECC values over the indicated population, Methods). Rug plots show mean attention weights for a set of 30 *Pvalb*-peak pairs (Methods) in `Pvalb` and `L5-ET-CTX` subclass populations. **b–c)** Attention weight for the same, co-present gene–peak pair can differ across cell populations due to systematic context differences. Welch’s t-test p-values are used to assess significance of such differences for gene–peak pairs that overall receive high attention weights (high attn pairs, *≥* 99.5^th^ percentile in (a)) that are also co-present in a pair of subclasses. A given gene can attend to the same peak more in `Pvalb` cells than in `L5-ET-CTX` cells (b) and vice versa (c). 9 such genes (e.g., *Srr* and *Myt1l*) appear with 23 distinct peaks (filled circles) above the significance threshold across panels (b) and (c). **d–e)** Same analysis, but for a pair of closely related supertypes `L5-ET-CTX-2` and `L5-ET-CTX-4`. Dashed lines in b–e indicate the Benjamini–Hochberg cut-off for significance (BH *q* = 0.05), and the numbers indicate the number crossing the cut-off and total number of comparisons in that plot. **f)** Pseudo-bulked chromatin accessibility values for the two subclasses and peaks in the *Pvalb*–peak pairs aligned to the genomic coordinate axis. While accessibility of this region is similar across the two subclasses, 15 of the 30 peaks receive significantly higher attention (dark blue color) within the `Pvalb` subclass. **g–h)** Cell-level distributions of attention weight (normalized such that uniform attention over the context is at 1.0) for the gene–peak pairs annotated in (b–c); asterisks indicate significance (BH *q ≤* 0.05); horizontal lines show group means.

Finally, we calculate the attention received by gene–peak pairs whose expression is restricted to a subset of subclasses. To account for unbalanced co-presence across subclasses, we use an information-theoretic metric, the effective cluster count (ECC).^44^ If a pair is co-expressed in only one subclass, its ECC value will be 1, whereas higher values reflect broader co-presence across multiple subclasses (**Methods**).

We find that median attention increases as gene–peak co-presence becomes more subclass-specific, i.e., at lower ECC thresholds: the median attentions for pairs with ECC less than 10, 2, and 1.1 are at the ∼86^th^, 87^th^, and 91^st^ percentiles of the background, respectively (**Figure 5a**). This suggests that Bloom attends to pairs of features based on their ability to distinguish cell identity and learns biologically meaningful attention values despite reporting on trillions of feature pairs (gene–gene, gene–peak).

### Bloom attends to feature pairs in a context-dependent manner

We investigate whether the cell context can produce significantly different attention values for given feature pairs between transcriptomically defined populations, when controlled for multiple testing. As a first case study, we explore the gene–peak pairs with average attention values at or above the 99.5^th^ percentile of the background (**Figure 5a**) and two classical cortical neuronal subclasses; excitatory neurons with extratelencephalic projections in layer 5 of cortex (`L5-ET-CTX`) and Parvalbumin-expressing cortical inhibitory neurons (`Pvalb`). **Figure 5b,c** shows the statistical significance of differences in mean attention between the two cell populations for every gene–peak pair co-present in both (Welch’s t-test; **Methods**). Between these two transcriptomically distant populations, fewer than 1,300 pairs are co-present (**Methods**), with roughly half receiving higher mean attention in `L5-ET-CTX` than in `Pvalb` (and vice versa). After correcting for multiple testing,^45^ we observe that 25% of `L5-ET-CTX` pairs and 32% of `Pvalb` pairs exhibit statistically significant differences in mean attention (*q <* 0.05), suggesting that Bloom can attend to feature pairs at distinct levels based on the cell context, despite its large library of genomic features.

We repeat this procedure on two supertypes, `L5-ET-CTX-2` and `L5-ET-CTX-4`, both of which are under the `L5-ET-CTX` subclass in the transcriptomic hierarchy.^4^ We observe that ∼10,000 feature pairs are co-present between these two closely related supertypes. Approximately 1% of these pairs are significantly differentially attended, suggesting that some of them may be useful for selective access to- or manipulation of these populations.

We next focus on a well-known marker gene, *Pvalb*. It is the classical marker gene for `Pvalb` neurons although it is also expressed in other populations, including `L5-ET-CTX` neurons (**Figure 5f**). Focusing on the 30 accessibility peaks around the *Pvalb* gene (**Methods**; **Figure 5f**), we observe distinct patterns of attention to *Pvalb*-peak pairs between the `L5-ET-CTX` and `Pvalb` cell populations (**Figures 5a,b,c**). Specifically, 15 of the 30 pairs are significantly more attended to in `Pvalb` cells than in `L5-ET-CTX` cells whereas no pairs are significantly more attended to in `L5-ET-CTX` cells (**Figure 5b,c,f**).

Finally, we observe the presence of genes that are preferentially associated with different peaks in the two populations. Here, we report on two example genes that were previously associated with neurological and psychiatric diseases: the *Srr* ^46, 47^–chr11:75,193,649 pair is attended to significantly more in `Pvalb` neurons (**Figure 5g**, left) compared to `L5-ET-CTX` neurons, and vice versa for the *Srr* –chr11:75,291,608 pair (**Figure 5g**, right). Similarly, the *Myt1l*^48, 49^–chr12:28,710,707 pair is attended to significantly more than the *Myt1l* –chr12:29,538,270 pair in `Pvalb` neurons (**Figure 5h**, left), and vice versa in `L5-ET-CTX` neurons (**Figure 5h**, right). Such preferential associations with different peaks can be useful for selectively manipulating a gene in a single subpopulation.

## Discussion

Cells in an organism differentiate to serve different roles. The mammalian brain is an extreme example with estimates suggesting thousands of cell types^4^ and spectacular displays of differentiation in morphology, connectivity, physiology, and genomic fronts. A key challenge is to define the cell state—a minimal and accurate description of its phenotype—and identify it consistently even though feature identities and counts may differ across experimental setups. To achieve this, here, we built a computational model that we refer to as Bloom. We focused on gene expression and chromatin accessibility characterizations as these can be profiled jointly with high throughput.^50^

During training, Bloom samples unbiased, random views of cells with a variable number of features. These views can choose features uniformly at random within an observation modality as well as across modalities (e.g., a set of measurements some of which profile gene expression while others profile peak accessibilities). Representations corresponding to different views of a cell are brought close to each other during training by the VISReg alignment objective, alongside training with the reconstruction objective. Indeed, we find that Bloom infers consistent representations for cells across gene expression and chromatin accessibility, and across different feature sets within each modality. Similarly, Bloom representations are relatively insensitive to the number of features used for cell views. We expect that this robustness will be useful in practice where different experimental setups characterize cells in different ways.

Bloom tokenizes (i.e., describes) genes and accessibility peaks by representations of the underlying DNA sequences, expanding upon models using protein embeddings for protein-coding genes only.^14, 15, 16^ We found that incorporating locations of these subsequences on the chromosome into the description further improves the quality of cell embeddings in Bloom. (A previous study on protein-coding genes only found no improvement on in-distribution data, SI Figure 8 in.^14^) As Bloom attends to relative locations of genomic features on the chromosome, it can infer feature interactions that might be obscure based on the sequence embeddings alone.

Cell representations learned by Bloom preserve the landscape of mouse brain cells substantially better than existing pre-trained models (**Figures 3** and **4**). Surprisingly, Bloom achieves this by training only on ∼300K cells. This is substantially smaller than the training corpora of those existing models. One explanation for this may be that Bloom’s cell views are unbiased and come from a dictionary of ∼30K genes and ∼1.7M peaks. This presents opportunities for data augmentation during training. We emphasize that Bloom’s training set was randomly sampled from a larger set, only ensuring that the training set is roughly balanced at the level of broad subclasses. Perhaps surprisingly, Bloom’s accuracy (as measured by linear classifiers) does not improve as more cells from the same population are added, either by increasing the per-subclass sampling cap or by adding unpaired RNA datasets. This is consistent with a lack of clear scaling relations between performance and training set size in tabular, single-cell datasets, unlike natural language corpora.^51^ Thus, building training sets by maximizing the variation represented in the corpus (e.g., including samples from organisms with different backgrounds, during development, aging, and disease) and not merely the cell count, should improve efficiency.

We studied the gene–peak associations inferred by Bloom via the weights of its first attention layer, which can act as a systematic hypothesis generation tool. We first showed that Bloom captures known enhancer associations. Perhaps more interestingly, the associations learned by Bloom do not merely reflect dataset-level correlations: a gene–peak pair that is highly attended in cells of one type may receive a significantly lower attention score in the presence of other genes that are frequently expressed (or peaks that are frequently accessible) in cells of a different type. We demonstrated this in case studies. Although these computationally inferred gene–peak associations should not be considered as true explanations before experimental validation,^52^ they provide targeted hypotheses (e.g., for selective manipulation) in a vast search space.

## Code availability

https://github.com/alleninstitute/bloom-biorxiv.

## Acknowledgments

We thank Marcus Hooper for help with data curation; Tyler Mollenkopf for help with project administration; Daniel Sebring, Daniel Schoonover, Aravind Neelakantan, Caleb Wilkinson, Rohit Thekkanal, and Alex Dinnouti for recommendations, setup, and management of compute resources on AWS; Arif Khan, Madhu Bussa, and Rick Buettner for AWS leadership support. We are appreciative of the financial and cloud infrastructure support from AWS’ Skilling and Social Impact and the Generative AI Innovation Center/Forward Deployed Engineers teams. This project was supported by Allen Institute. RG, IC, YM, RL, NG, SO, SC, ZY, MH, SM, HZ, US thank the Allen Institute founders, P. G. Allen and J. Allen, for their vision, encouragement and support.

## Methods

### Overview

The overall flow of data and the model architecture are shown in **Figures 2a,b**. Let *G*_*c*_ denote the set of genes and *P*_*c*_ denote the set of peaks for which measurements are made in a given cell *c*. Define the set of features *F*_*c*_ = *G*_*c*_ ∪ *P*_*c*_. Each feature in *F*_*c*_ (indexed by *f*) consists of a sequence encoding *s*_*f*_ , a type encoding *t*_*f*_ , a position encoding *r*_*f*_ , and the measured value in cell *c, x*_*fc*_. The sequence and position encodings are detailed in a subsequent section.

A *view v* of cell *c* is a subset of sampled features 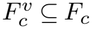 During training, a single entry of the mini-batch consists of two input views and a query view for a given cell. We use *v* ∈ {I, II} to denote the input views and *v* = Q to denote the query view.

Both input views are sampled from the same feature set, chosen at random from 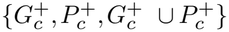, where 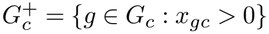 and 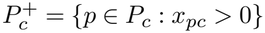 denote the non-zero gene and peak sets for a given cell. The input views consist of a variable number of features per view, 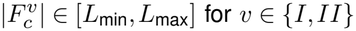, where *L*_min_ is a hyperparameter and *L*_max_ is the maximum input context length. The views are sampled such that a fraction *α* of 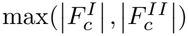 features is shared across the views. *α* is a hyperparameter that follows a schedule during training. While the query view 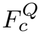 can be any subset of *F*_*c*_, here we set it to a fixed set of ∼8,000 differentially expressed genes to focus on features informative about brain cell types.

For each input view *v* ∈ {I, II}, the encoder maps 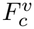 to a corresponding cell representation, 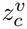, which is then routed to task-specific heads during model training. We consider reconstruction of genes and peaks (if present) in the query view as separate tasks. For these tasks, decoders use 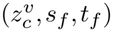 to predict 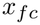 for features in 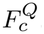. We also use a self-supervised learning task with the VISReg^25^ objective applied to 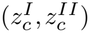 to learn view-invariant representations while preventing representation collapse. The task heads are discarded after model training. During inference, a single view of the cell is sampled from 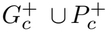 as specified for the particular evaluation to obtain a representation for that cell.

### Sequence and position encodings

The universe of genomic features including 32,129 genes and 1,747,701 chromatin accessibility peaks is defined using the mm10 (GRCm38) mouse reference genome. Each gene is referenced by a 500-bp interval centered at the transcription start site (TSS) of its longest transcript. Each peak is defined as a fixed 500-bp genomic interval based on an independent analysis of aggregated chromatin accessibility data from mouse brain cells. A fixed-length, context-aware sequence encoding (*s*_*f*_) is obtained for each feature with a genomic language model (gLM) such as Evo 2^21^ or AlphaGenome.^22^

Such gLMs provide a vector representation for each position in the input DNA sequence. To avoid potential edge effects and provide the gLM with additional context, the input sequence to the gLM for each feature is an expanded 2,048-bp window centered on the feature. We then mean-pool the representations for the positions corresponding strictly to the central 500-bp subsequence. We use the `blocks.28.mlp.l3` layer of the `evo2_7b` model checkpoint for Evo 2, and the PyTorch port^53^ with the `model_all_folds` checkpoint for AlphaGenome. The resulting 4,096-dimensional sequence encodings from Evo 2 and the 1,536-dimensional embeddings from AlphaGenome are subsequently projected to 768 dimensions using PCA and independently z-score normalized across each dimension before being used as input to Bloom.

The midpoint of a genomic feature is considered to be its location. Thus, each feature is associated with a chromosome identifier *ζ* and a coordinate *b* as per the reference genome. Analogous to the approach in Vaswani et al.,^6^ we represent position on a chromosome using *M* logarithmically spaced wavelengths *λ* between 100 bp and 100 Mbp as

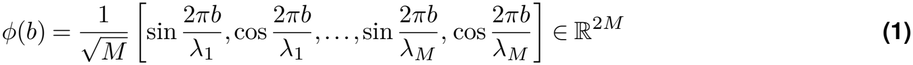

This ensures translational invariance of the inner product, i.e., 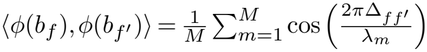 is a function of the coordinate difference Δ_*ff*_*′* = *b*_*f*_ − *b*_*f*_*′* . We also want to discourage interactions between features on different chromosomes. This can be achieved in expectation by defining the position encoding for the features as 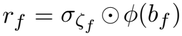, where *σ*_*ζ*_ ∈ {−1, +1}^2*M*^ is a fixed random sign vector drawn independently for each chromosome and ⊙ denotes the element-wise product. We use 128-dimensional position encodings with *M* = 64 wavelengths in [10^2^, 10^8^].

### Bloom encoder

The encoder consists of a single multi-head attention layer that operates on the input features, followed by a Perceiver block.^24^ The Perceiver block includes a multi-head cross-attention layer with learnable latents, which reduces the context length for subsequent self-attention layers to a fixed, small number (relative to the input context length). The output of the final layer of the Perceiver block is mean-pooled to obtain the cell representation.

The sequence encodings and genomic location provide distinct information about the input features. Further, it is the relative position that can retain consistent meaning across species. In a given self-attention head, the attention weights between a pair of features are computed with an inner product between the query and key vectors.^6^

We define sequence heads so that the query, key, and value vectors for an input feature explicitly use the sequence encodings. Accordingly, for each sequence head we use 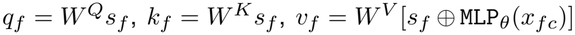 where *W* ^*Q*^, *W* ^*K*^, *W* ^*V*^ are learned parameters, MLP_*θ*_ is a multi-layer perceptron with parameters *θ* and ⊕ denotes vector concatenation.

We define position heads to leverage the construction of positional encodings at length scales of interest for each head. For position head *h*, the query, key, and value vectors are 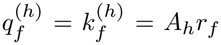 and 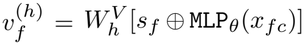. Here, *A*_*h*_ = diag(*a*_*h*1_, *a*_*h*1_, … , *a*_*hM*_ , *a*_*hM*_) is a fixed matrix with non-negative *a*_*hm*_ applied to position encoding components corresponding to wavelength *λ*_*m*_, and 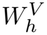 are learned parameters. In particular, *a*_*hm*_ ∝ exp(−(ln *λ*_*m*_ − ln Λ_*h*_)^2^*/*2*ρ*^2^) and is normalized so that 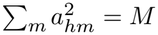. This sets the scale for the inner product in **Equation 1**. Bandwidth *ρ* and Λ_*h*_ are hyperparameters chosen to specify length scales over which genomic feature interactions are prioritized in a given position head. We set *ρ* = 1.15 ≈ ln(10)*/*2, and use eight position heads with Λ_*h*_ logarithmically spaced in [10^3^, 10^7^]. Note that inter-chromosomal interactions between genomic features are permitted through the sequence heads as well as standard attention heads.

### Model configuration and training

In this section we provide the key configuration parameters for the model used in all of the experiments shown in **Figure 2d** onwards. Within the encoder, the input attention layer consists of 8 sequence heads, 8 position heads, and 4 standard attention heads, each with a 32-dimensional output per feature per head. We used Evo 2 sequence encodings as described above to train the model unless mentioned otherwise. The cross-attention layer of the Perceiver block uses 256 learned latents, each of which has 1,024 dimensions. This is followed by 8 (latent) self-attention layers with 8 heads per layer, with *d*_model_ set to 1,024. The feed-forward blocks use GELU activations and the Perceiver block follows the pre-LN architecture,^54^ with dropout of 0.1 throughout the model. The latents at the final layer are mean-pooled and layer-normalized to obtain the 1,024-dimensional cell representation.

Query embeddings incorporate sequence and feature, modality, and batch metadata, together with library-size information. Modality-specific Bernoulli decoders are MLPs that factorize detection probability into biological, library-depth, and technical components, each parameterized by a sigmoid output. For the self-supervised learning objective, the embeddings are first projected with an MLP to 256 dimensions, with batch normalization and GELU activations. The invariance term of the VISReg objective matches this projection across different views of the same cell. The regularization terms use 256 random projections (called sketches) to make this representation isotropic and Gaussian.

The model was implemented in PyTorch, and distributed training was achieved with PyTorch Lightning DDP. Model training took 10 days on 2× p5e nodes with 8 NVIDIA H200 GPUs per node. We used bfloat16 mixed precision for a smaller memory footprint, which enabled larger batch sizes and context lengths. The batch size per GPU was set to 2 views of 64 cells, leading to a global batch of 1,024 cells. The maximum context length for each of the input as well as query views was set to 8,192. The backward pass was computed at the end of every step (i.e., without gradient accumulation). We used AdamW with *β*_1_ = 0.9, *β*_2_ = 0.999, *ϵ* = 10^*−*8^, and weight decay 0.01, excluding biases and normalization parameters from weight decay. Gradients were clipped to an L2 norm of 1.0. The learning rate increased linearly from 10^*−*8^ to 10^*−*4^ over the first 1,000 steps and was subsequently held constant.

### Datasets

#### Training

The training corpora for experiments in **Figure 2f** are documented in **Table 1**. The model version used for evaluations in **Figures 2-5** was trained with the `1k-mult` dataset, except for the relevant experiments in **Figure 2f**. per subclass (min. 500).

#### Evaluation

Held-out subsets of the training data, as well as other datasets used for evaluations throughout the manuscript, are detailed in **Table 2**. These datasets capture a diversity of single-cell resolution genomic measurement technologies for mammalian brain tissue.

**Table 2.** Held-out evaluation datasets used to assess single-cell representation quality. ^†^Multiome data from the Allen Institute (unpublished). Both human multi-region datasets (Siletti and SEA-AD) span the same eight cortical and hippocampal regions of interest (ROIs): FI (frontoinsular cortex), HIP (hippocampus), ITG (inferior temporal gyrus), LEC (lateral entorhinal cortex), MEC (medial entorhinal cortex), MTG (middle temporal gyrus), STG (superior temporal gyrus), and V1C (primary visual cortex).

| Dataset | Species | *n* (cells) | *n* (types) | Notes |
|---|---|---|---|---|
| Multiome (a)<sup>†</sup> | Mouse | 15,750 | 315 subclasses | Whole brain (76 dissection regions); paired snRNA-seq and snATAC-seq; 50 cells per subclass. |
| Multiome (b)<sup>†</sup> | Mouse | 36,090 | 1,203 supertypes | Whole brain (76 dissection regions); paired snRNA-seq and snATAC-seq; 30 cells per supertype. |
| scRNA-seq<sup>4</sup> | Mouse | 15,500 | 310 subclasses | Whole brain (89 dissection regions); 10x v2 and v3; 50 cells per subclass. |
| snRNA-seq (Macosko)<sup>34</sup> | Mouse | 14,850 | 297 subclasses | Whole brain; single-nucleus 10x v3; 50 cells per subclass. |
| VISp Patch-seq<sup>35, 36</sup> | Mouse | 6,599 | 18 subclasses | Primary visual cortex, layers 1–6b (98.9% of cells); 4,715 GABAergic and 1,884 glutamatergic cells; 10–1,952 cells per subclass. |
| Aging snRNA-seq<sup>37</sup> | Mouse | 13,560 | 226 supertypes | Adult and aged (6,780 cells each; 30 per supertype per age group); isocortex, hippocampal formation, cerebral nuclei, hypothalamus, midbrain and pons. |
| snRNA-seq (Siletti)<sup>38</sup> | Human | 22,634 | 66 subclasses | 8 cortical and hippocampal ROIs; 3 donors; 30–800 cells per subclass. |
| AD snRNA-seq (SEA-AD)<sup>39</sup> | Human | 65,417 | 143 supertypes | Same 8 ROIs; 32 donors aged 68–100 (14 with dementia, 18 without; Braak 0 and II–VI); 10x v3.1 and Multiome (RNA only); 11–3,646 cells per supertype. |

**Table 3.** Architecture comparison of the single-cell models used for evaluations in this study. Parameter counts only list trainable parameters.

| Model | Parameters | Layers | Heads | *d*<sub>model</sub> | Training context |
|---|---|---|---|---|---|
| Bloom | 131M | 10 | 20 | 1024 | 8,192 |
| SE-600M | 603M | 16 | 16 | 2048 | 2,048 |
| TF-Metazoa | 444M | 12 | 16 | 2048 | 2,047 |
| TF-Exemplar | 542M | 12 | 16 | 2048 | 2,047 |
| UCE | 675M | 33 | 20 | 1280 | 1,024 |

### Community models

There are a large number of transformer-based models for unimodal, single-cell transcriptomic data. Here we chose prominent, recent models trained on large datasets to compare the performance of Bloom on unimodal tasks. While UCE^14^ and Transcriptformer^15^ are trained on multi-species data, SE-600M^16^ was trained only on human data. To facilitate comparisons with SE-600M we restricted the gene universe to 15,732 strict 1:1 Ensembl mouse orthologs for human protein-coding genes.

Efforts to model chromatin accessibility with a similar class of models are comparatively less developed. Multimodal or even unimodal chromatin accessibility datasets are not as comprehensive in terms of species and tissue systems at single-cell resolution. The feature universe is often species- and tissue-specific, which makes direct comparisons with community models such as Epi-Agent,^55^ SCARF,^56^ CLM-X+^57^ (human only), Nephrobase+^58^ (kidney) and Atacformer^59^ (peripheral blood mononuclear cells) difficult to interpret.

### Evaluation tasks

#### Classification

Linear probing is a proxy method to evaluate the quality of representations via linear decoding of labels of interest.^60^ Representations are obtained from a frozen model and used to train a downstream linear classifier. Here we use a stratified K-fold cross-validation setup with K=10 and the scikit-learn implementation of logistic regression (default inverse regularization strength C=1.0) applied to standardized representations unless mentioned otherwise. In the genes-only setting, we use the *n* most highly expressed genes per cell to obtain the cell representation. In the peaks-only setting, we use a random sample of 8,192 non-zero peaks per cell to obtain the cell representation.

Classification performance is reported with the adjusted balanced accuracy as implemented in scikit-learn 1.9.0 (`balanced_accuracy_score` with `adjusted=True`), with error bars denoting ±1 standard error of the mean (SEM) across cross-validation folds where relevant. In practice, these values tend to be similar to Macro F1 score, which is another commonly used metric to quantify classification performance.

For the experiments in **Figure 2d**, we obtain cell representations for *n* ∈ {500, 1,000, 2,000, 4,000, 8,000}. The classifier reuse procedure for this experiment is described next. A classifier is trained on held-out cells using representations obtained with *n* = 8,000. This classifier is then applied to cells in the validation set using representations obtained for the target *n* value.

#### Clustering

The clustering task probes whether the embedding organizes cells of the same type into contiguous regions of latent space without supervision. We run *k*-means on the embedding using the `sklearn.cluster.KMeans` implementation with *n*_init_=10 random restarts, taking *k* equal to the number of unique classes in `obs`[`class_level`]. Reported metrics are adjusted Rand index (ARI), adjusted mutual information (AMI), and normalized mutual information (NMI), each computed between the k-means assignments and the ground-truth labels.

#### Label transfer

The label-transfer task evaluates cross-dataset, cross-modality, and cross-species generalization by training a classifier on embeddings from a source dataset and evaluating it on a disjoint target dataset. We use the same approach as in the classification task, except that the logistic regression is fit once (no cross-validation) on the entire source dataset. Evaluation is restricted to the class labels shared between the source and target sets.

#### Batch effect

The batch-effect task quantifies how well an embedding integrates sources that differ in a nuisance covariate (dataset origin, sequencing modality) while preserving biological signal. We use the scIB benchmark suite^61^ via `scib_metrics` and compute batch-correction scores against a reference label. Metrics for batch-correction scores include graph iLISI, local mixing of batch labels, kBET acceptance rate and label-aware kBET.

### Attention-based analyses

The sequence heads of the model could be used to generate hypotheses about gene–peak interactions. Here we focus on genes as queries, and other genes or peaks in the input context as keys within the sequence heads. The average attention over all the sequence heads for a given gene–peak or gene–gene pair is referred to as the attention weight for all analyses in **Figure 5**.

#### Background pairs

The input to the model consists of randomly drawn non-zero genes and peaks per cell. Pairs constructed from each such gene and peak combination are considered as background gene–peak pairs. Observations (*n* = 962B pairs) across a full pass over the training set `1k-mult` are used to construct the background distribution.

#### Proximal cis pairs

Any peak on the same chromosome as a given gene and within a distance of 1 Mbp from its TSS is considered a proximal cis peak for that gene. These are further restricted to a set that is co-present in at least 50 cells across the training dataset (`1k-mult`), leading to 23,715,315 gene–peak pairs.

#### Enhancer pairs

We used a published list of enhancers^43^ as a starting point. This list defines variable-length (min: 250 bp; max: 2,202 bp) genomic intervals (mm10) that are experimentally found to be strong and on-target enhancers among cortical subclasses. The list also identifies nearby gene(s) on the same chromosome that have highly correlated expression patterns in relation to enhancer accessibility in a pseudo-bulked setting. We match these enhancers to the universe of peaks and genes used in this study as a first step, and keep only peaks for which the overlap with the enhancer is at least 250 bp. The 127 unique gene–peak pairs for which the primary target subclass is available in our training data are used for attention-based analysis.

#### *Pvalb*–peak pairs

The *Pvalb* gene is expressed in cell populations of both the `Pvalb` and `L5-ET-CTX` subclasses. A 36-kbp interval containing the *Pvalb* gene TSS has a similar chromatin accessibility pattern across the two subclasses, **Figure 5f**. The accessibility values in **Figure 5f** are normalized by the mean signal in the 1-kbp window centered on the *Pvalb* TSS for each track. We consider all 30 peaks in our universe that fall in this interval and pair each with the *Pvalb* gene to define the *Pvalb*–peak pairs.

#### ECC

The exponential of Shannon entropy is the Hill number of order 1, and is commonly used as a diversity index.^44^ To account for differences in sample size across groups, we compute a rarefied effective cluster count, `ECC`, defined as the expectation of this quantity over random draws of fixed size *n*.

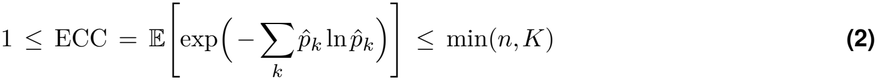

In particular, we draw *n* cells from the pool of cells for which a given gene–peak pair is co-present, uniformly at random without replacement. We determine the empirical distribution 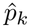 as the fraction of drawn cells that belong to subclass *k*. The total number of subclasses is denoted as *K*. `ECC` is 1 when all drawn cells belong to one subclass, and the upper bound corresponds to the case where each cell in the draw belongs to a different subclass.

#### Context-specific attention

Particular gene–peak pairs are sampled in the context of genes and peaks in a given cell, and changing the context can shift the attention assigned to that pair. We use this to look for gene–peak pairs that shift significantly across cell populations of interest. Here we consider cell types at two distinct levels of the taxonomy: distant subclasses, `060 Pvalb_Gaba` and `023 L5-ET-CTX_Glut`, and related supertypes `0088 L5-ET-CTX_Glut 2` and `0090 L5-ET-CTX_Glut 4`. We refer to these subclasses as `Pvalb` and `L5-ET-CTX`, respectively, and the supertypes as `L5-ET-CTX-2` and `L5-ET-CTX-4`, respectively, everywhere else for convenience. We restrict this search to proximal cis gene–peak pairs that receive high attention scores overall as per **Figure 5a** (≥ 99.5^th^ percentile). For a comparison across populations, the gene–peak pair of interest must be non-zero (i.e., co-present) in both populations, which filters out many gene–peak candidates. To these candidates we also add the *Pvalb*–peak pairs. For each pair, we independently sample contexts from the full pool of non-zero genes and peaks and calculate the mean attention score over multiple draws from the same cell. The context is filled up to 4,096 entries each time. We draw 5 contexts per cell for up to 80 cells per population in which the gene and the peak are co-present. The resulting distribution of context-averaged attention weights per cell is scaled so that uniform attention over the context is 1. We call this the normalized attention weight and use Welch’s t-test to determine if these distributions have different means across the populations. This is a two-sided test, and the Benjamini–Hochberg correction is performed once for all pairs in the candidate set. We split the candidates based on the direction of the mean difference (**Figure 5g–h**) to highlight cases where the normalized attention weight changes in opposite directions for different peaks attended to by the same gene, depending on the cell type context.

